# SARS-CoV-2 Omicron is specifically restricted in its replication in human lung tissue, compared to other variants of concern

**DOI:** 10.1101/2022.03.31.486531

**Authors:** Or Alfi, Marah Hamdan, Ori Wald, Arkadi Yakirevitch, Ori Wandel, Esther Oiknine-Djian, Ben Gvili, Hadas Knoller, Noa Rozendorn, Hadar Golan, Sheera Adar, Olesya Vorontsov, Michal Mandelboim, Zichria Zakay-Rones, Menachem Oberbaum, Amos Panet, Dana G. Wolf

## Abstract

SARS-CoV-2 Omicron variant has been characterized by decreased clinical severity, raising the question of whether early variant-specific interactions within the mucosal surfaces of the respiratory tract could mediate its attenuated pathogenicity. Here, we employed *ex vivo* infection of native human nasal and lung tissues to investigate the local-mucosal susceptibility and innate immune response to Omicron, compared to Delta and earlier SARS-CoV-2 variants of concern (VOC). We show that the replication of Omicron in lung tissues is highly restricted compared to other VOC, whereas it remains relatively unchanged in nasal tissues. Mechanistically, Omicron induced a much stronger antiviral interferon response in infected tissues compared to Delta and earlier VOC - a difference which was most striking in the lung tissues, where the innate immune response to all other SARS-CoV-2 VOC was blunted. Our data provide new insights to the reduced lung involvement and clinical severity of Omicron.

## INTRODUCTION

The recently evolved SARS-CoV-2 Omicron variant has been shown to exhibit increased transmissibility and escape from humoral immunity generated by previous infections and vaccines ^1–3^. Importantly, accumulating clinical-epidemiological observations have demonstrated that Omicron is associated with milder disease compared with Delta and earlier variants of concern (VOC) ^2,4,5^. The decreased clinical severity of Omicron has been partly attributed to the presence of pre-existing population immunity ^2^. Additionally, it has been proposed that intrinsic viral factors could play a part in its milder disease course. This notion was suggested by studies in animal models showing that Omicron infection caused milder lung pathology ^6–8^, and by the reported inefficient replication of Omicron in human alveolar organoids and *ex vivo* infected lung tissues ^9,10^. It has been shown that Omicron enters cells by a different route than other variants and does not spread as efficiently by fusion, thereby limiting viral infection in the lungs, where cell-fusion plays a role in viral transmission^2,9–12^. These studies highlight the multifactorial yet incompletely resolved mechanisms underlying the decreased pathogenicity of Omicron.

The innate immune response is known to play a key-role in SARS-CoV-2 infection severity^13–15^. We therefore reasoned that the early innate immune responses to Omicron within the respiratory tract could potentially mediate its milder clinical severity. To investigate the local-mucosal susceptibility and response to Omicron we have used our recently established *ex vivo* SARS-CoV-2 infection models in native 3D human nasal and lung tissues, which recapitulate viral infection in the upper and lower respiratory tract^16^. We identified distinctive patterns of susceptibility and antiviral interferon responses to Omicron, compared to Delta and precedent VOC, which were most remarkable in lung tissues, and provide new clues to the reduced clinical severity of Omicron.

## RESULTS

### SARS-CoV-2 Omicron exhibits restricted replication in human lung tissues

Nasal and lung tissues maintained viable as integral organ cultures as described^16^, were infected in parallel with Omicron and Delta, using the same viral inoculum. Viral replication kinetics were monitored between 2-72 hours post infection by quantitative measurements of tissue-associated viral sub-genomic (sg)-mRNA and secreted infectious virus progeny, as described^16^. To control for the expected tissue-to-tissue variations (reflecting the natural diversity of different donors) we used five and four independent lung and nasal tissues, respectively. Whereas Omicron and Delta demonstrated similar replication kinetics in the nasal tissues (Figure 1A), the replication of Omicron in the lung tissues was highly restricted compared to the productive replication of Delta (Figure 1B). This relative replication restriction, which was most apparent at late times post infection, was confirmed by confocal microscopy analysis, showing the near-absence of infected cells in Omicron-infected lung tissues (Figure 1C). The Delta replication kinetics in the lung tissues was overall similar to those of D614G, Alpha, and Beta variants (Figure 1-figure supplement 1). Hence, the restricted replication of Omicron in the lung tissues distinguished it form all precedent SARS-CoV-2 VOC examined.

**Figure 1.**
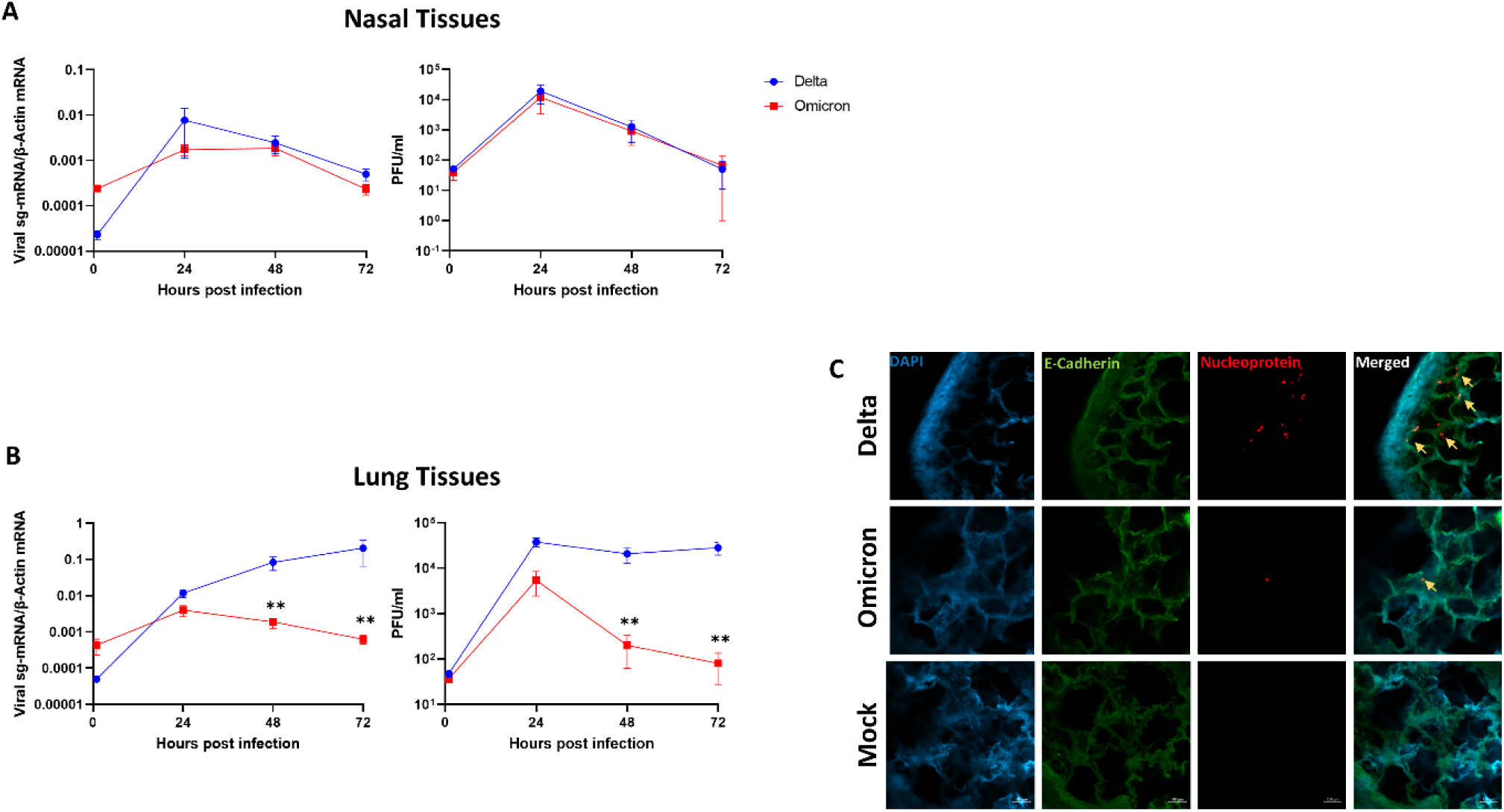
SARS-CoV-2 Omicron and Delta replication kinetics in human nasal and lung tissues. Nasal (**A**) and lung (**B**) organ cultures were (each) infected in parallel with Omicron and Delta (10^5^ PFU/well). The levels of tissue-associated viral subgenomic (sg)-mRNA (left panels), and infectious virus progeny released from the same infected tissues (right panels) represent mean values (± SEM) of four and five independent nasal and lung tissues, respectively, each tested in four biological replicates. (**C**) Representative confocal micrographs of whole-mount lung tissues at 72h post infection. **, p<0.01. Statistics were performed using multiple paired, two-tailed Student’s t test.

### Omicron elicits enhanced antiviral interferon response in human respiratory tissues

We have previously shown that SARS-CoV-2 infection (the ancestral isolate USA-WA1/2020) induced a robust innate immune response in nasal tissues, albeit a highly restricted innate immune response in lung tissues^16^. To compare the innate immune response triggered by Omicron versus Delta, we examined the expression of interferons (IFNs) and representative antiviral interferon stimulated genes (ISG) in lung tissues upon parallel infection with the two variants. The selected ISG included MX1, IFI6, ISG15, and IFIT1, which have been previously demonstrated to exhibit broad-acting antiviral activities and to inhibit SARS-CoV-2 replication^17^. Interestingly, employing RT-qPCR, we showed that Omicron infection elicited a vigorous lung-tissue innate immune response, with substantial induction of the expression of INFλ and ISGs (Figure 2A). We could not detect upregulation of INFα and INFβ in the infected tissues (data not shown). The strong interferon response induced by Omicron was in sharp contrast to the low ISG response of the same lung tissues to Delta. We also showed, in parallel infection experiments, that the restricted lung-tissue response to Delta was common to all other VOC tested, including D614G, Alpha, and Beta (whereas the same lung tissues still exhibited a strong response to influenza virus) (Figure 2-figure supplement 1). In line with the enhanced interferon response triggered by Omicron in the lung tissues, Omicron also induced some enhancement of the interferon response in infected nasal tissues, compared with Delta (Figure 2 - figure supplement 2). Yet, it was notable that the nasal tissues (unlike the lung tissues) already exhibited a strong response to Delta (Figure 2 - figure supplement 2), which was generally consistent with our previous observations^16^. Thus, the enhancement of the innate immune response to Omicron was most remarkable in the lung tissues, where the response to all other SARS-CoV-2 VOC was largely restricted.

**Figure 2.**
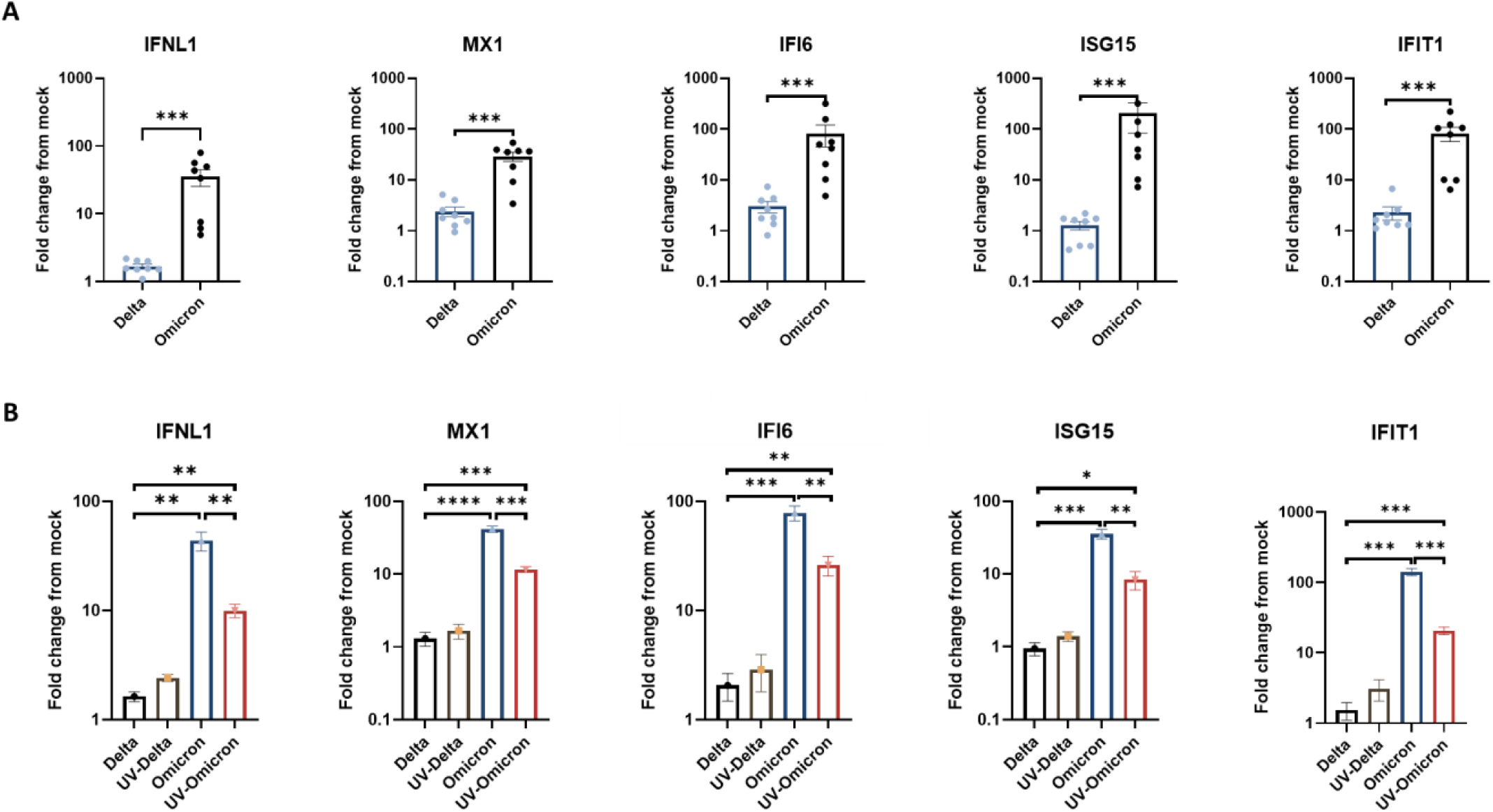
Lung tissue interferon response to SARS-CoV-2 Omicron and Delta. Lung organ cultures were infected in parallel with Omicron and Delta (10^5^ PFU/well), and the effect of infection on the expression of the indicated innate immunity genes, measured by RT-qPCR at 24h post infection, is presented as fold-change from mock-infection. (**A**) the mean values (± SEM) in 8 independent tissues, each tested in 4 biological replicates. (**B**) the mean values (± SEM) in a representative tissue (tested in 4 biological replicates) exposed in parallel to infectious versus UV-inactivated viruses. The results shown in panel B represent at least three independent tissues. *, *P* < 0.05; **, *P* < 0.01; ***, *P* < 0.001; ****, *P <* 0.0001

To further define whether the relative enhancement of the innate response to Omicron in the lung tissues was dependent on active viral replication, we compared the induction of ISG following exposure to infectious versus UV-inactivated Omicron and Delta virions. surprisingly, a significant induction of ISG was observed following exposure of the lung tissues to UV-inactivated Omicron (but not Delta) virions (Figure 2B), despite the absence of de-novo viral gene expression (Figure 2 - figure supplement 3). This finding indicated that the enhanced lung response to Omicron is already triggered, at least in part, by virion structural component/s upon initial virus-cell contact or entry, preceding viral gene expression.

## DISCUSSION

Omicron infection has been characterized by decreased clinical severity, raising the question of whether early variant-specific interactions within the mucosal surfaces of the respiratory tract could mediate its attenuated pathogenicity. Using *ex vivo* infection models of native human lung and nasal tissues, we show that the replication competence of Omicron in lung tissues is highly restricted compared to Delta and precedent VOC, whereas it remains relatively unchanged in nasal tissues. The susceptibility of the nasal viral entry site to Omicron may support person-to-person transmission, whereas its restricted replication in the lungs could contribute to the milder clinical course of Omicron.

Our findings reveal a new mechanism, whereby the significantly enhanced antiviral interferon responses to Omicron compared to earlier VOC, which was most striking in the lung tissues (where the innate immune response to all other SARS-CoV-2 VOC was blunted), could limit its pathogenicity. Innate immune defenses have been shown to play a crucial role in the control of SARS-CoV-2 infection, and impaired local interferon responses in the respiratory tract have been associated with severe COVID-19^13,14,17^. Hence, an augmented mucosal innate immune response to Omicron could capture the virus at the upper respiratory tract and limit viral infection and pathology in the lungs. In addition to the upregulated ISG, the observed induction of IFNλ by Omicron in the lung tissues (Figure 2) is noteworthy, given the reported key-role of mucosal IFN-III in the protection against life-threatening SARS-CoV-2^17^, as well as in the prevention of excessive inflammatory damage^18^. Our finding that the interferon response to Omicron was already triggered upon initial viral attachment/entry further implies a causative relation between the enhanced early antiviral response and the restricted spread of Omicron in lung tissues. This latter finding may be related to the predominant use of the endocytic entry pathway by Omicron (as opposed to Delta)^2,9,11,12^, which could lead to early activation of unique endosomal Toll-like receptors. The mechanism by which Omicron triggers the enhanced interferon response in human respiratory tissues remains to be elucidated.

Our study has several limitations. Native respiratory tissues in organ culture are relatively short-lived (up to 7 days in culture). Thus, our *ex vivo* infection models mirror early events of infection and do not address the late phase of viral transmission or the combined effects of the local and systemic immune responses. Nonetheless, our studies recapitulate SARS-CoV-2 infection and innate immune response within the authentic multicellular complexity of both the upper and the lower human respiratory tract, containing tissue-specific compositions of cell types, including immune cells, and extracellular matrix. To date, we are not aware of studies that have examined the distinctive innate tissue responses to Omicron as related to its altered replication phenotype in the lower respiratory tract. Such data are critical to better understand and address the evolution of SARS-CoV-2 into a less virulent human-tropic virus. In summary, our studies in native human nasal and lung tissues infected *ex vivo* reveal a significantly enhanced interferon response to Omicron, compared to precedent SARS-CoV-2 VOC. The findings imply that the early induction of antiviral ISG, which was most prominent in lung tissues, could play a part in the restricted replication and pathology of Omicron in the lungs. They provide insights for the attenuated pathogenicity of Omicron, and for further studies of pathways involved in the enhanced mucosal innate immune responses to this newly evolving variant.

## MATERIALS AND METHODS

### Cells and viruses

Simian kidney Vero E6 (ATCC CRL-1586), Calu-3 (ATCC HTB-55), Madin-Darby Canine Kidney (MDCK, ATCC CCL-34™) cells and H1299-ACE2 overexpressed cells (kindly provided by Dr. Alex Sigal)^1^ were maintained in Dulbecco’s Modified Eagle Medium (DMEM; Biological Industries, Beit Haemek, Israel), supplemented with 10% fetal bovine serum, 2 mM L-Glutamine, 10 IU/ml Penicillin, and 10 μg/ml streptomycin (Biological Industries, Beit Haemek, Israel). An early pandemic SARS-CoV-2 D614G isolate (GISAID ID: EPI_ISL_10125580), an Alpha B.1.1.7 isolate (GISAID ID: EPI_ISL_10125211), a Delta, B.1.617.2 isolate (GISAID ID: EPI_ISL_9837720) and an Omicron B.1.1.529 isolate (GISAID ID: EPI_ISL_7869197) were isolated from positive nasopharyngeal swab samples. The Beta variant B.1.351 (GISAID ID: EPI_ISL_678615) was generously provided by Dr. Alex Sigal. All viruses were isolated and propagated (2 passages) in Calu-3 cells, and sequence verified. Influenza virus A(H1N1) pdm09 (NIBRG-121xp, Cat# 09/268; obtained from NIBSC, UK) was propagated in MDCK cells. The virus titers of cleared infected cells- and tissue supernatants were determined by a standard plaque assay on H1299-ACE2 cells (SARS-CoV-2) or MDCK cells (influenza virus).

### Preparation and infection of nasal turbinate and lung organ cultures

Nasal turbinate and lung organ cultures were prepared and infected as previously described^16^. In brief, inferior nasal turbinate tissues were obtained from consented patients undergoing turbinectomy procedures, and lung tissues (the tumor free margins) were obtained from consented patients undergoing lobectomy operations. The study was approved by the Hadassah Medical Center (#0296-20-HMO) and the Sheba Medical Center (#2832-15-SMC) Institutional Review Boards. Fresh tissues were kept on ice until further processed at the same day. The tissues were sectioned by a microtome (McIlwain Tissue Chopper; Ted Pella, INC.) into thin slices (250 μm-thick slices; each encompassing ∼10 cell layers), and incubated in 0.3 ml of enriched RPMI medium (Biological Industries, Beit Haemek, Israel) (for the nasal turbinate tissues) or DMEM/F-12 medium (Biological Industries, Beit Haemek, Israel) with MEM Vitamin Solution (Biological Industries, Beit Haemek, Israel) (for the lung tissues), with 10% fetal bovine serum, 2.5 μg glucose/ml, 2 mM glutamine, 10 IU/ml penicillin, 10 μg/ml streptomycin, and 0.25 μg/ml amphotericin B, at 37°C, 5% CO2. The tissues were processed and infected at the same day (the day of harvesting; Day 0). For infection of the organ cultures, the tissues were placed in 48-well plates and inoculated with the respective virus (1×10^5^ PFU/well in 0.3 ml) for 12h to allow effective viral adsorption. For UV-inactivated virus exposure experiments in the lung tissues, the medium containing 1×10^5^ PFU in 0.3 ml was pre-exposed to UV for 1 hour and validated for complete loss of infectivity. Following viral adsorption, the cultures were washed three times (in 0.3 ml of complete medium) and further incubated for the duration of the experiment, with replacement of the culture medium every 2 to 3 days. Tissue viability was monitored by the mitochondrial dehydrogenase enzyme (MTT) assay as previously described^16^. All infection and tissue processing experiments were performed in a BSL-3 facility.

### Whole-mount tissue immunofluorescence

Tissues were fixed in 4% formaldehyde for 24hours, washed in PBS and transferred to 80% ethanol. The tissues were permeabilized by 0.3% Triton-X100 in PBS (PBST) and further incubated with Animal-Free Blocker® (Vector laboratories, Cat# SP-5035-100) to block nonspecific antibody binding, followed by incubation with the primary antibodies in Animal-Free Blocker® at room temperature overnight. The tissues were than washed 4 times in PBST, incubated with the secondary antibodies in Animal-Free Blocker® for at room temperature overnight, washed 4 times with PBST, and incubated with 4’,6-diamidino-2-phenylindole (DAPI, 10uM, Abcam, Cat# ab228549) as a nuclear stain. The following primary antibodies were used: α-E-Cadherin (Mouse monoclonal, 1:100, Abcam, ab1416; for the detection of epithelial cells), α-SARS-CoV-2 Nucleocapsid (Rabbit monoclonal, 1:500, Abcam, ab271180). The following secondary antibodies were used: Donkey anti-Mouse IgG pre-adsorbed, Alexa Fluor® 568 (1:250, Abcam, Cat# ab175700), Goat anti-Rabbit IgG Highly Cross-Adsorbed Alexa Fluor Plus 647 (1:250, Thermo Fisher Scientific, Cat# A32733). For tissue clearing, stained preparations were dehydrated with 100% Ethanol for 1h, and later submerged and mounted in ethyl-cinnamate (99%; Sigma, Cat# 112372) as previously described ^16^. Whole-mount tissues were visualized using a Nikon A1R confocal microscope and were analyzed using NIS Elements software (Nikon).

### RNA purification and quantification

Infected- and mock-infected organ cultures and the respective supernatants were flash-frozen and stored at -80°C until assayed. RNA was extracted using NucleoSpin RNA Mini kit for RNA purification (Macherey-Nagel, Cat #740955.250) according to the manufacturer’s instructions, and subjected to reverse transcription, using High-Capacity cDNA Reverse Transcription Kit (Thermo Fisher Scientific, Cat#). Quantitative real time (RT)-PCR was performed on a Quantstudio 3™ (Thermo Fisher Scientific) instrument, using Fast SYBR™ Green Master Mix (Thermo Fisher Scientific, Cat# 4385614), or TaqMan™ Fast Advanced Master Mix (Thermo Fisher Scientific, Cat# 4444558). The employed primers and probe sequences are listed in Supplementary Table 1.

### Statistical analysis

All data, presented as means ± standard errors of the mean (SEM), were analyzed using paired, two-tailed t test in GraphPad Prism 9 software (GraphPad Software Inc., San Diego CA). P values of <0.05 were considered significant.

## Data availability

All the sequences of the virus isolates used are available in GISAID. The accession numbers of viral sequences used in this study are listed above. The raw data for all graphs presented in this paper are included as source data.

## ACKNOWLEGEMENTS

The study was supported by the Israel Science Foundation (530/18 & 1728/20), the Israel Ministry of Science & Technology (3-16964), and the Hadassah France Association.

## FIGURE SUPPLEMENTS

**Figure 1-figure supplement 1.**
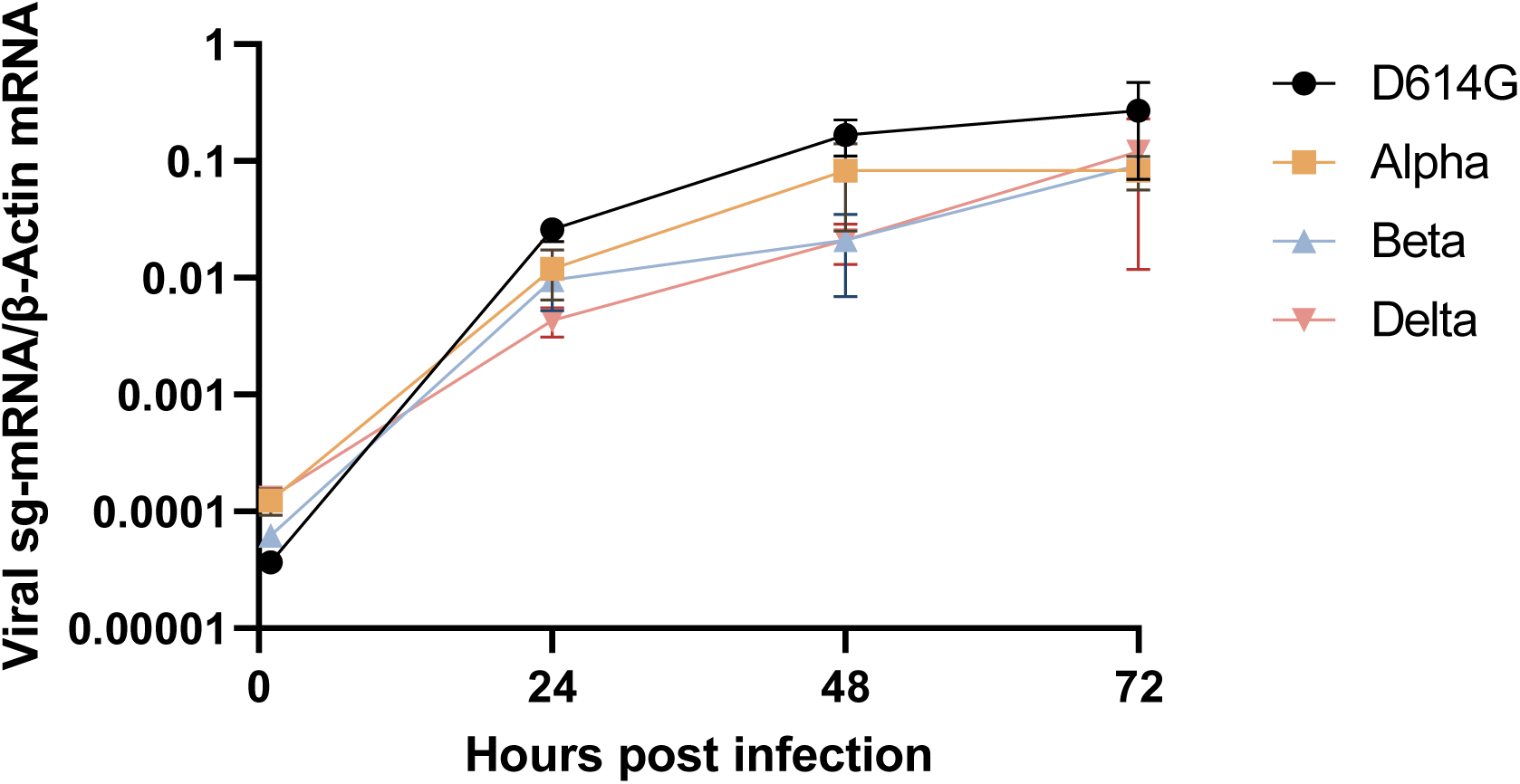
Replication kinetics of SARS-CoV-2 variants in human lung tissues. Lung organ cultures were infected in parallel with the indicated variants. Levels of tissue-associated SARS-CoV-2 N gene subgenomic (sg)-mRNA were determined by RT-qPCR and normalized to β-actin. The data shown represent the mean values (± SEM) of at least three independent tissues, each tested in 4 biological replicates.

**Figure 2-figure supplement 1.**
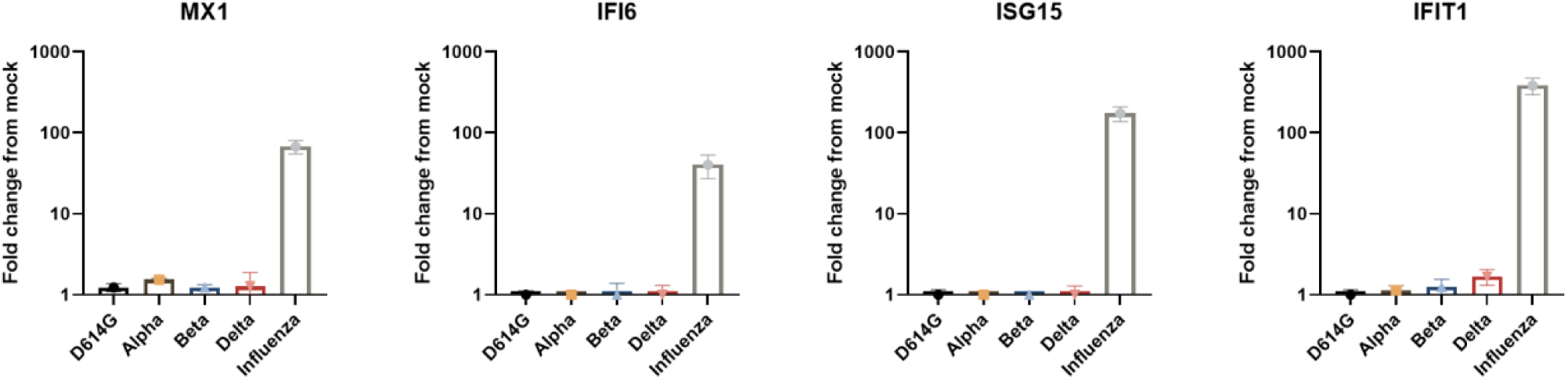
Lung tissue ISG response to SARS-CoV-2 variants and influenza. Lung organ cultures were infected in parallel with the indicated variants, and with Influenza A/H1N1 (10^5^ PFU/well). RNA was extracted from mock- and infected tissues at 24h post infection, and the effect of infection by the indicated viruses on the expression of interferon-stimulated genes (ISG) is presented as fold-change from mock infection. The data shown represent the mean values (± SEM) of at least three independent tissues, each tested in 4 biological replicates.

**Figure 2-figure supplement 2.**
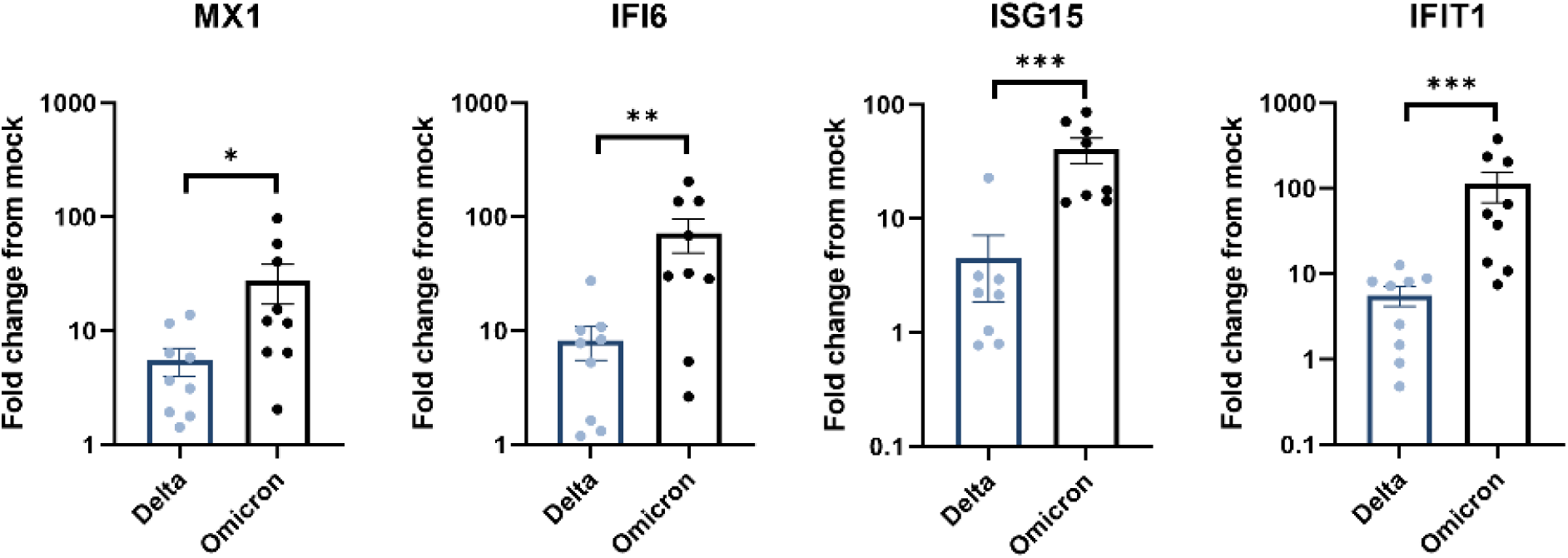
Nasal tissue ISG response to SARS-CoV-2 Omicron and Delta. Nasal organ cultures were infected in parallel with Omicron and Delta (10^5^ PFU/well), and the effect of infection on the expression of the indicated interferon-stimulated genes (ISG), measured by RT-qPCR at 24h post infection, is presented as fold-change from mock-infection. The data shown represent the mean values (± SEM) of 9 independent tissues, each tested in 4 biological replicates. *, *P* < 0.05; **, *P* < 0.01; ***, *P* < 0.001

**Figure 2-figure supplement 3.**
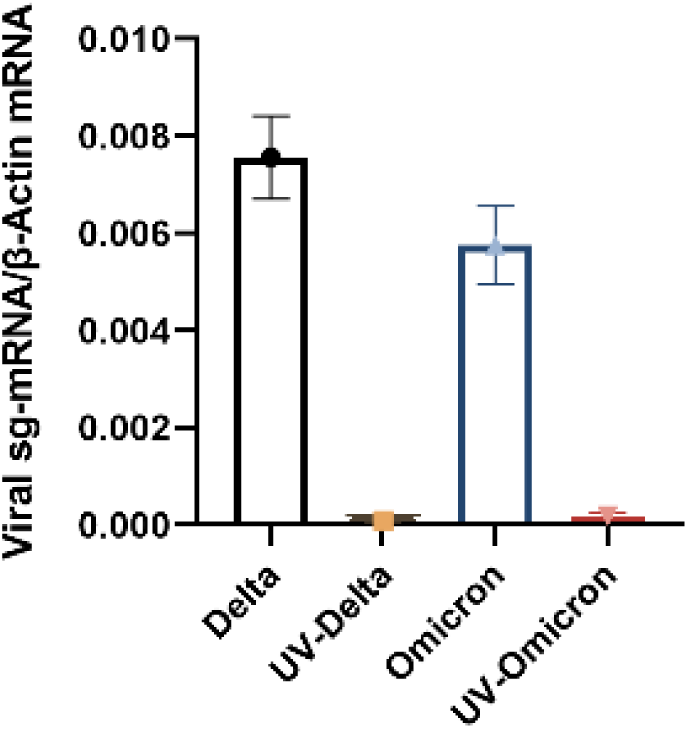
Viral subgenomic (sg)-mRNA in lung tissues exposed to infectious versus UV-inactivated SARS-CoV-2 Omicron and Delta. Lung organ cultures were exposed in parallel to infectious versus UV-inactivated Omicron and Delta variants (10^5^ PFU/well), and the levels of tissue-associated viral sg-mRNA were measured by RT-qPCR at 24h post infection. The results shown as mean values (± SEM) in a representative tissue (tested in 4 biological replicates), represent at least three independent lung tissues.

